# Data-driven subtyping of executive function related behavioural problems in children

**DOI:** 10.1101/158949

**Authors:** Joe Bathelt, Joni Holmes, the CALM team, Duncan E. Astle

**Affiliations:** MRC Cognition and Brain Sciences Unit, University of Cambridge, 15 Chaucer Road, Cambridge CB2 7EF

## Abstract

**Introduction:** Many developmental disorders are associated with deficits in controlling and regulating behaviour. These difficulties are frequently observed across multiple groups of children including children with diagnoses of attention deficit hyperactivity disorder (ADHD), specific learning difficulties, autistic spectrum disorder, or conduct disorder. The co-occurrence of these behavioural problems across disorders typically leads to comorbid diagnoses and can complicate intervention approaches. An alternative to classifying children on the basis of specific diagnostic criteria is to use a data-driven grouping that identifies dimensions of behaviour that meaningfully distinguish groups of children.

**Methods:** The sample consisted of 442 children identified by health and educational professionals as having difficulties in attention, learning and/or memory. The current study applied community clustering, a data-driven clustering algorithm, to group children by similarities across scales on a commonly used rating scale, the Conners-3 questionnaire. Further, the current study investigated if the groups identified by the data-driven algorithm could be identified by white matter connectivity using a structural connectomics approach combined with partial least squares analysis.

**Results:** The data-driven clustering yielded three distinct groups of children with symptoms of either: (1) elevated inattention and hyperactivity/impulsivity, and poor executive function, (2) learning problems, and (3) aggressive behaviour and problems with peer relationships. These groups were associated with significant inter-individual variation in white matter connectivity of the prefrontal and anterior cingulate cortex.

**Conclusion:** In sum, data-driven classification of executive function difficulties identifies stable groups of children, provides a good account of inter-individual differences, and aligns closely with underlying neurobiological substrates.

## Introduction

Research into behavioural problems is often guided by clinical definitions of disorders. These diagnostic definitions can pose significant challenges for understanding the nature of behavioural problems and uncovering neurobiological mechanisms. A case in point is attention deficit hyperactivity disorder (ADHD), which is commonly diagnosed on the basis of elevated symptoms of inattention and hyperactivity/impulsivity ^1^. While these symptoms are prominent features of ADHD, they are also common among other frequently diagnosed developmental disorders including dyslexia^2^, autism^3^ and conduct disorder^4^. The presentation of these behavioural problems can vary very widely across children with the same diagnosis ^5^. In practical terms, this often results in children receiving comorbid diagnoses or diagnoses that do not fully capture the broad and complex range of difficulties they face. As a result, standard intervention approaches tailored for one specific diagnosis (e.g. medication for ADHD) do not meet children’s complex individual needs. The overlap in symptom presentation between, and the variability within, disorders complicates research into the causes and treatment of common developmental difficulties. The aim of the current study was to use a data-driven approach to identify groups of children with similar dimensions of behavioural problems and to investigate the relationship between white matter connectivity and these groupings.

Traditional categorical diagnostic approaches to understanding developmental disorders (e.g. ICD-10, World Health Organization, 1992) have considerable practical advantages by facilitating clinical decision making ^6^. However, current diagnoses are based on a tradition of clinical insight rather than knowledge about pathophysiological mechanisms ^7^. The slow progress in understanding the mechanisms leading to neurodevelopmental disorders, and associated difficulties with identifying effective treatments, may in large part be attributable to the heterogeneous aetiology within traditional diagnostic classes. Subgroups within diagnostic categories may follow different pathways with a superficially similar clinical presentation ^8^. For instance, early-onset and later-onset ADHD form distinct categories that are associated with different aetiological mechanisms and different comorbidities ^9^.

An alternative to classifying children on the basis of specific diagnostic criteria is to use a data-driven grouping that identifies similarities in behaviour that meaningfully distinguish groups of children. Recent evidence from taxonomic and heritability studies support this approach to understanding ADHD ^10,11^. A dimensional approach can provide the practical advantages of clearly defining categories, while also identifying the most pertinent behavioural characteristics for investigations into pathophysiological mechanisms. The current study uses a data-driven community clustering approach to group children by the similarity of their behavioural problems. In contrast to widely employed factor analytic approaches that aim to capture *measured variables* through a smaller set of latent factors related to theoretical constructs (e.g. grouping questionnaire items that relate to hyperactivity or inattentivity), the clustering approach adopted in this study *groups individual children* by similarities in their behavioural ratings. This alternative approach is made possible by recent advances in network science methods, which we apply to a large dataset comprised of children identified as having problems in attention, learning and/or memory, who were referred to the study by educational and clinical professionals working in various specialist children’s services. This large sample includes children with specific, multiple and no diagnoses. The sample is therefore not already restricted to children who have met particular diagnostic criteria, but instead consists of a broad range of children who are struggling educationally. This offers the opportunity to identify the most pertinent behavioural dimensions, while side-stepping the biases inherent in recruiting according to current diagnostic classification.

Data-driven approaches to identifying behavioural profiles are not straightforward. Most clustering algorithms necessitate *a priori* assumptions, like the geometrical properties of the cluster shape, the tuning of some parameters, or setting the number of desired clusters. These assumptions are difficult to make, but network science provides a possible solution. This is the study of complex networks, which represent relationships between data as a network of nodes connected by edges. This methodological approach provides a mathematical tool for quantifying the organisation of networks and the relationships between the nodes within them ^12^. Defining subdivisions of highly-connected nodes within a network, so called communities, is an area of network science that has received considerable attention as it applies to many real world problems ^13^. In the case of psychometric data, the network can represent the similarity of scores between participants. Community detection makes it possible to define subgroups of participants that are most similar, while being as distinct as possible from other subgroups. The aim of the current study was to identify clusters in a large sample of children referred for cognitive and learning difficulties by the similarity of their executive function-related behavioural problems using a community detection approach based on parent ratings on the Conner’s questionnaire. This scale is routinely administered in health care and educational settings, and in many clinics in the U.K. it is used to measure behavioural problems at home and school to aid in the diagnosis of ADHD.

One of the aims of data-driven nosology is to identify behavioural dimensions that are more closely related to biological mechanisms. In the current study, we explored differences in white-matter connectivity between the groups identified through the community detection. White matter maturation is a crucial process of brain development that extends into the third decade of life ^14^, and which relates closely to cognitive development ^15-17^. It is thought to support cognitive development through better communication and integration between brain regions, particularly over longer distances. Accordingly, the brain can be modelled as a network of brain regions connected by white matter, commonly referred to as a connectome. Brain regions vary in the number of their connections – their node degree -which gives an indication of their importance for the network ^19, 20^,. To explore which brain regions were most closely linked to the behavioural profiles identified through consensus clustering, we used a multi-variate dimension reduction technique called partial least squares (PLS) ^21^. In our analysis, PLS defined brain components that maximally distinguished the behaviourally defined groups.

## Participants and Methods

### Participants

The sample consisted of ratings on 442 children, using the Conner’s Parent Rating Short Form 3^rd^ edition ^22^, referred to as “Conners 3” from here on. The ratings were completed by parents or caregivers (age: mean=110.51 months; SE=1.24; Range=62-215; 295 male) as part of a larger ongoing study at the Centre for Attention, Learning and Memory (CALM) at the MRC Cognition and Brain Sciences Unit, University of Cambridge. Children were recruited to the CALM research clinic on the basis of having problems in attention, learning and / or memory that had come to the attention of a professional working in schools (e.g. special needs teacher) or specialist children’s community services (e.g. clinical or educational psychologists, speech and language therapists or paediatricians). During the clinic visit, children completed a wide range of cognitive assessments while their parents/caregivers filled in questionnaires about the child’s behaviour. Children were also invited for an MRI structural scan (see Figure 1 for attainment). The data reported here include three questionnaires and the MRI data. Exclusion criteria for referrals were significant or severe known neurological disorders, problems in vision or hearing that were uncorrected, or having a native language other than English. This study was carried out in accordance with the Declaration of Helsinki and was approved by the local NHS research ethics committee (Reference: 13/EE/0157). Written parental/caregiver consent was obtained and children provided verbal assent.

**Figure 1:**
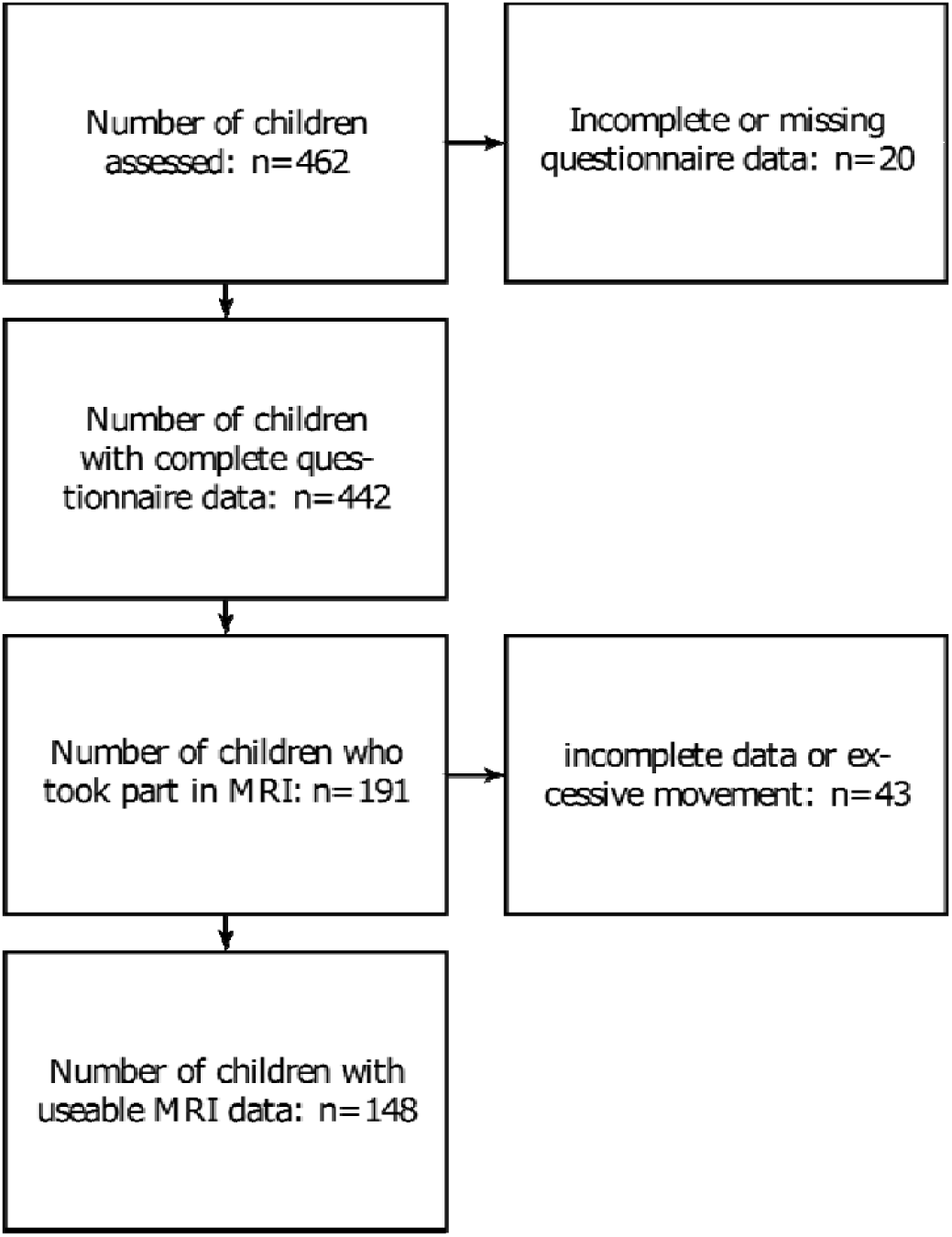
Overview of data included in behavioural and connectome analysis

Some children in the broad sample of children referred for problems relating to attention, learning, and/or memory, had received diagnoses through standard community services (see Table 1 for a breakdown of diagnoses). Among the children with a diagnosis, ADHD was the most common. Other diagnostic labels were rare. Therefore, diagnostic labels were grouped together for the comparison between diagnoses and the data-driven groups. Primary diagnoses of dyslexia, dyscalculia, or dysgraphia were summarised as ‘learning deficits’. Primary diagnoses of autism spectrum disorder, autism, or Asperger syndrome were summarised as ‘ASD’. Other labels, like OCD, depression, anxiety, or developmental delay occurred only in a few individuals and were grouped as ‘other’.

**Table 1:**
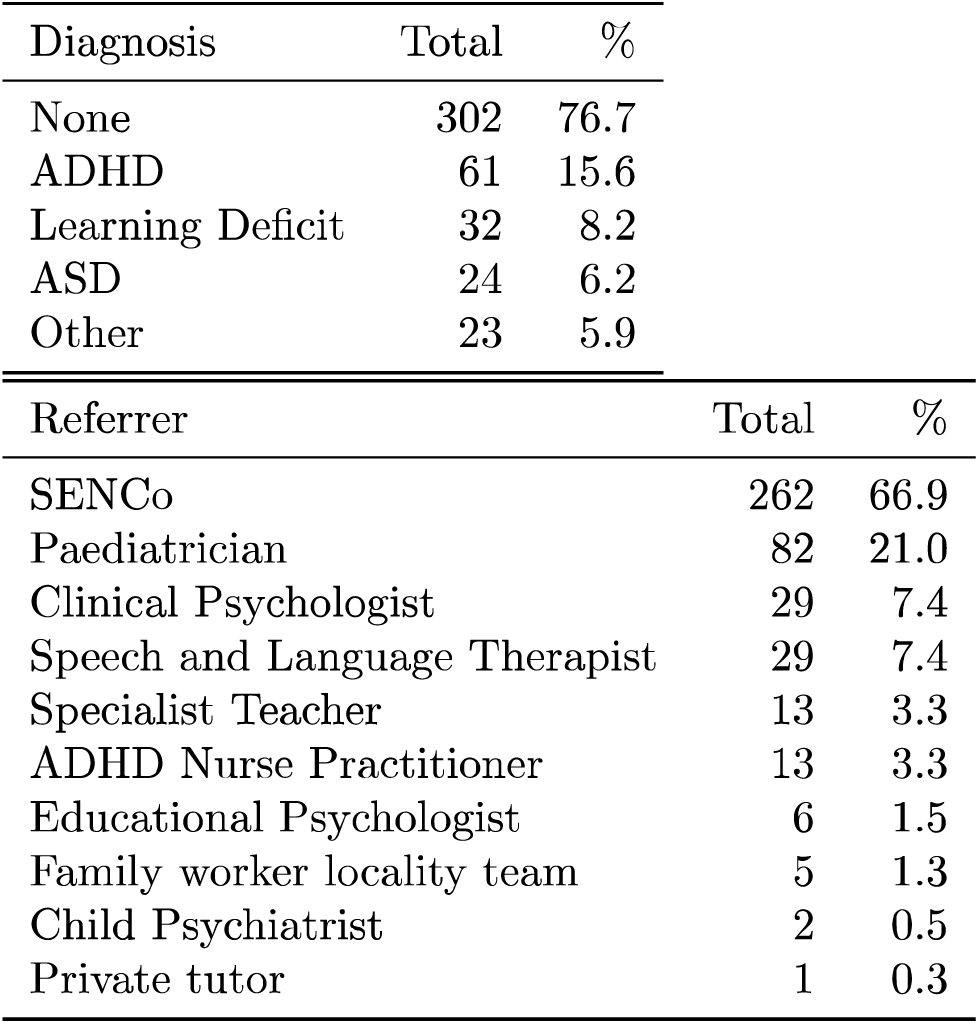
Breakdown of children by pre-existing diagnoses and referral routes. Abbreviations: ADHD: attention deficit hyperactivity disorder, ASD: autism spectrum disorder, SENCo: special educational needs coordinator

| Diagnosis | Total | % |
| --- | --- | --- |
| None | 302 | 76.7 |
| ADHD | 61 | 15.6 |
| Learning Deficit | 32 | 8.2 |
| ASD | 24 | 6.2 |
| Other | 23 | 5.9 |

| Referrer | Total | % |
| --- | --- | --- |
| SENCo | 262 | 66.9 |
| Paediatrician | 82 | 21.0 |
| Clinical Psychologist | 29 | 7.4 |
| Speech and Language Therapist | 29 | 7.4 |
| Specialist Teacher | 13 | 3.3 |
| ADHD Nurse Practitioner | 13 | 3.3 |
| Educational Psychologist | 6 | 1.5 |
| Family worker locality team | 5 | 1.3 |
| Child Psychiatrist | 2 | 0.5 |
| Private tutor | 1 | 0.3 |

## Behavioural Analysis

### Questionnaire Data

#### Conners-3

The Conners scale ^22^ is a parent questionnaire designed to assess behavioural difficulties associated with ADHD and related disorders. It is well validated with good psychometric integrity (Internal consistency: Cronbach’s alpha=0.91 [Range: 0.85-0.94]; Factorial validity: RMSEA=0.07 based on confirmatory factor analysis in a replication sample; for details see Conners, 2013). Questionnaire items are summarised into six subscales (Inattention, Hyperactivity/Inattention, Learning Problems, Executive Function, Aggression, Peer Problems) and a total ADHD score is also derived. T scores of 60 and above are indicative of clinical levels of problems. A high proportion of children in the sample had scores in this range on each of the subscales (see Table 2).

**Table 2:** Scores on each scale of the Conners-3 questionnaire (inattention, hyperactivity/impulsivity, learning problems, executive function, aggression, peer relationships) for the entire sample. The last two columns indicate the total number and the percentage of children in the sample with T scores in the clinical range on each scale. Abbreviations: std= standard deviation of the mean.

|  | mean | std | min | max | *T* >60 | *T* >60% |
| --- | --- | --- | --- | --- | --- | --- |
| Inattention | 79.74 | 11.955 | 40 | 90 | 398 | 90.0 |
| Hyperactivity/Impulsivity | 72.87 | 16.388 | 40 | 90 | 315 | 71.3 |
| Learning Problems | 75.95 | 11.912 | 42 | 90 | 390 | 88.2 |
| Executive Function | 73.81 | 12.906 | 40 | 90 | 363 | 82.1 |
| Aggression | 62.95 | 17.268 | 34 | 90 | 205 | 46.4 |
| Peer Relationships | 71.94 | 17.973 | 44 | 90 | 290 | 65.6 |

The Conners also contains two validity scales to detect response bias, i.e. the rater tries to convey an overly positive or negative impression to secure a certain outcome ^22^. The validity scales indicated a possibly overly negative response style for 80 responses. Highly negative scores may indicate extreme problems in the rating domains or a negative bias of the rater, which may overestimate the child’s difficulties. Analyses were carried out including and excluding ratings with high Negative Impression scores.

The Behavioral Rating Inventory of Executive Function (BRIEF) is a questionnaire about behaviours associated with executive function problems for parents of children aged 5 to 18 years ^23^. There are eight subscales measuring behaviour problems related to inhibition, shifting, emotional control, initiation, working memory, planning/organising, organisation of materials, and monitoring.

The Strengths and Difficulties Questionnaire (SDQ) is a parent-rated scale for children aged 8 to 16 years. It provides ratings for emotional symptoms and prosocial behaviour as well as scores for problems related to behavioural conduct, hyperactivity/inattention, and peer relationships.

## Community Detection

Community detection is an optimisation clustering method. Networks in the current analysis represented the child-by-child correlations across the 6 scales of the Conners 3 questionnaire. The community algorithm starts with each network node, i.e. child, in a separate community and then iteratively parcellates the network into communities to increase the quality index (Q) until a maximum is reached. The current study used the algorithm described by Rubinov and Sporns ^19^ as implemented in the Brain Connectivity Toolbox (https://sites.google.com/site/bctnet/) version of August 2016. This algorithm is not deterministic and may yield different solutions at each run. In order to reach a stable community assignment, we applied the consensus clustering method described by Lancichinetti and Fortunato ^24^. In short, an average community assignment over 100 iterations was generated. The community assignment was then repeated further until the community assignment did not change between successive iterations. The analysis was implemented in Python 2.7.11. The code for the entire analysis is available online (http://www.github.com/joebathelt/).

## Statistical Analysis

Groups defined by the community detection algorithm were compared on scales of the Conners 3 questionnaire. Shapiro-Wilk tests indicated that scores within groups deviated from normality assumptions ^25^. Group contrasts were therefore based on non-parametric Mann-Whitney *U* tests. The Bonferroni method was used to account for multiple comparisons. Statistical tests were carried out using Scientific Python (SciPy) version 0.17.0 implementation ^26^.

## Structural connectome

The aim of this analysis was to explore whether the data-driven grouping was related to differences in brain structure. To this end, white matter connectivity of brain regions was estimated from diffusion-weighted images. Next, we employed a multivariate, dimension-reduction technique to relate the white-matter connectivity of brain regions to the group assignment.

## Participant sample for the connectome analyses

A subset of 191 families agreed to the neuroimaging part of the study. A total of 43 scans were excluded for poor quality, i.e. incomplete scan data, visually identified movement artefact, maximum displacement in the diffusion sequence above 3mm as determined by FSL eddy (see Figure 1 for an overview of attrition). The final sample consisted of 148 complete datasets (behaviour, T1, dwi). The MRI sample did not significantly differ in age from the behavioural sample (MRI sample [months]: mean=117.05, std=27.436, t(359)=1.34, p=0.181). The ratio of groups defined in the analysis of the behavioural sample was similar in the MRI subsample (MRI sample: C1: 0.36, C2: 0.33, C3: 0.30).

## MRI data acquisition

Magnetic resonance imaging data were acquired at the MRC Cognition and Brain Sciences Unit, Cambridge U.K. All scans were obtained on the Siemens 3 T Tim Trio system (Siemens Healthcare, Erlangen, Germany), using a 32-channel quadrature head coil. The imaging protocol consisted of two sequences: T1-weighted MRI and a diffusion-weighted sequence.

T1-weighted volume scans were acquired using a whole brain coverage 3D Magnetisation Prepared Rapid Acquisition Gradient Echo (MP RAGE) sequence acquired using 1mm isometric image resolution. Echo time was 2.98 ms, and repetition time was 2250 ms.

Diffusion scans were acquired using echo-planar diffusion-weighted images with an isotropic set of 60 non-collinear directions, using a weighting factor of b=1000s*mm ^-2^, interleaved with a T2-weighted (b = 0) volume. Whole brain coverage was obtained with 60 contiguous axial slices and isometric image resolution of 2mm. Echo time was 90 ms and repetition time was 8400 ms.

## Structural connectome construction

The white-matter connectome reconstruction followed the general procedure of estimating the most probable white matter connections for each individual and then obtaining measures of fractional anisotropy (FA) between regions (see Figure 2). The details of the procedure are described in the following paragraphs. In the current study, MRI scans were converted from the native DICOM to compressed NIfTI-1 format using the dcm2nii tool (http://www.mccauslandcenter.sc.edu/mricro/mricron/dcm2nii.html). Subsequently, a brain mask was derived from the b0-weighted volume of the diffusion-weighted sequence and the entire sequence was submitted for correction for participant movement and eddy current distortions through FSL’s eddy tool. Next, non-local means de-noising ^27^ was applied using the Diffusion Imaging in Python (DiPy) v0.11 package ^28^ to boost signal to noise ratio. The diffusion tensor model was fitted to the pre-processed images to derive maps of fractional anisotropy (FA) using dtifit from the FMRIB Software Library (FSL) v.5.0.6 ^29^. A spherical constrained deconvolution (CSD) model ^30^ was fitted to the 60-gradient-direction diffusion-weighted images using a maximum harmonic order of 8 using DiPy. An alternative analysis with a constant solid angle (CSA) model is present in the Supplementary Materials section. Next, probabilistic whole-brain tractography was performed based on the CSD model with 8 seeds in any voxel with a General FA value higher than 0.1. The step size was set to 0.5 and the maximum number of crossing fibres per voxel to 2.

**Figure 2:**
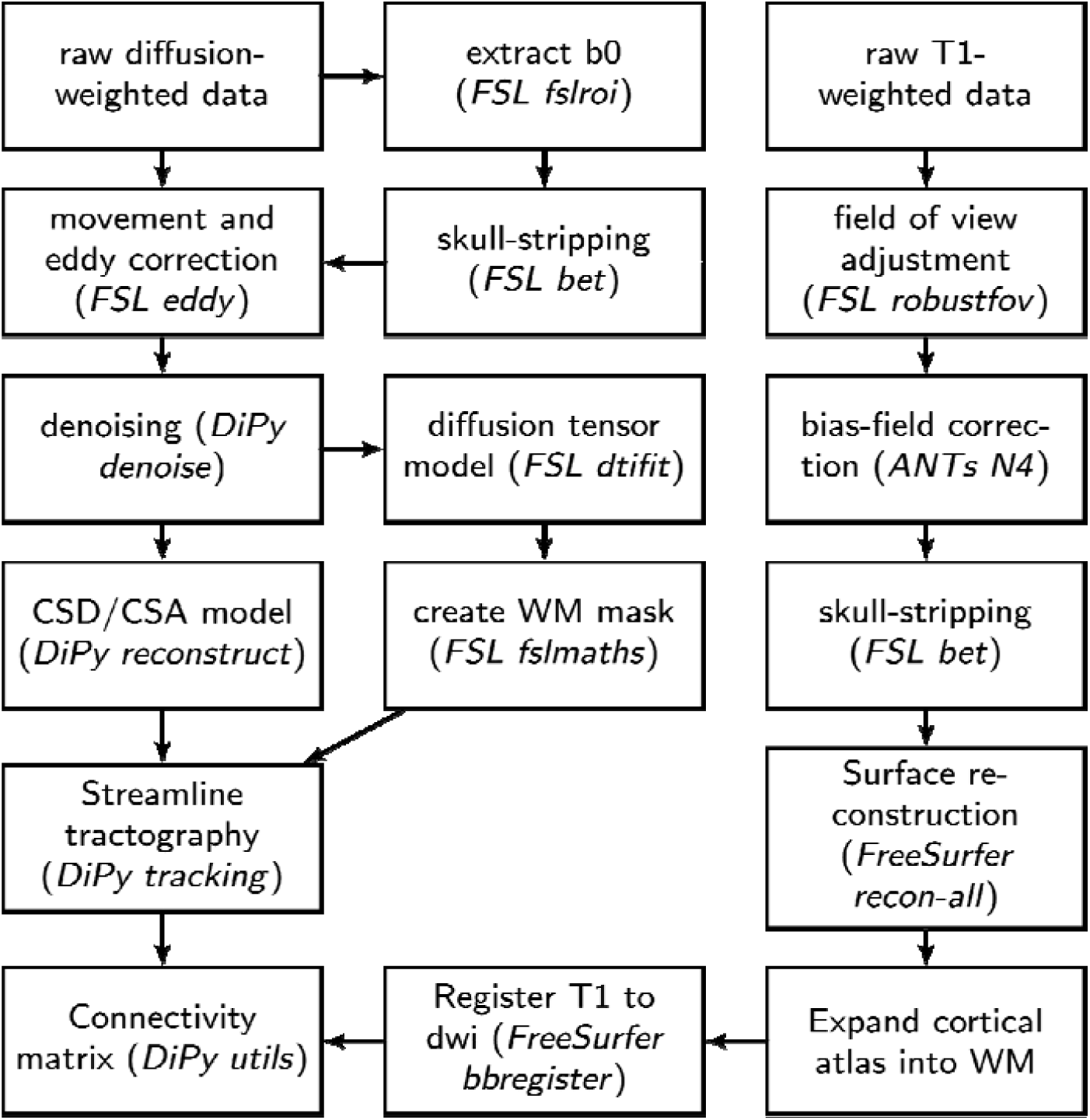
Overview of processing steps for structural connectome estimation

For ROI definition, T1-weighted images were pre-processed by adjusting the field of view using FSL’s robustfov, non-local means denoising in DiPy, deriving a robust brain mask using the brain extraction algorithm of the Advanced Normalization Tools (ANTs) v1.9 ^31^, and submitting the images to recon-all pipeline in FreeSurfer v5.3 (http://surfer.nmr.mgh.harvard.edu). Regions of interests (ROIs) were based on the Desikan-Killiany parcellation of the MNI template ^32^ with 34 cortical ROIs per hemisphere and 17 subcortical ROIs (brain stem, and bilateral cerebellum, thalamus, caudate, putamen, pallidum, hippocampus, amygdala, nucleus accumbens). The surface parcellation of the cortex was transformed to a volume using the aparc2aseg tool in FreeSurfer. Further, the cortical parcellation was expanded by 2mm into the subcortical white matter using in-house software. In order to move the parcellation into diffusion space, a transformation based on the T1-weighted volume and the b0-weighted image of the diffusion sequence was calculated using FreeSurfer’s bbregister and applied to volume parcellation.

For each pairwise combination of ROIs, the number of streamlines intersecting both ROIs was estimated and transformed to a density map. A symmetric intersection was used, i.e. streamlines starting and ending in each ROI were averaged.

The weight of the connection matrices was based on fractional anisotropy (FA). To obtain FA-weighted matrices, the streamline density maps were binarized after thresholding and multiplied with the FA map and averaged over voxels to obtain the FA value corresponding to the connection between the ROIs. This procedure was implemented in-house based on DiPy v0.11 functions ^28^. False positive streamline can introduce spurious results in structural connectome analyses ^33^. To remove spurious connections in the FA-weighted and streamline density-weighted networks, consensus thresholding was applied so that only connections that were present in more than 60% of the sample were retained ^34^.

## Statistical analysis of connectome data

For the analysis of the connectome data, the node degree of each node in the network wass calculated for each participant. Partial least squares regression was used to identify thee linear combination of brain areas that best explained group membership for the groupss identified through community clustering. The PLS model was evaluated by fitting the modell to random selection of 60% of the data and evaluating the model fit in a test set of 40%. The root-mean-squared error of a model based on the training data was significantly lowerr when assessed with the test data compared to randomly shuffled samples (10-fold cross-validated RMSE: mean=0.35, SE=0.025; permuted sample: mean=0.81, SE=0.018;; permutation test: *p*=0.002).

The contribution of brain regions to the PLS latent variables was evaluated in a bootstrapp procedure in which 60% of the sample was randomly selected and the PLS model wass fitted (1000 permutations). The loading of brain regions onto PLS latent variables was expressed as the mean loading divided by the standard error across permutations. AA Procrustes rotation was applied to align the factor across iterations of the permutationn procedure. All procedures were implemented using scikit-learn functions v0.18.1 under Python v2.7.12 ^35^.

## Results

### Community Detection indicates three subgroups

The current study employed graph theory to derive clusters of children with similar profiles across ratings on the Conners-3 questionnaire. The community detection algorithm in conjunction with consensus clustering arrived at a stable solution with three clusters. The quality index (Q=0.55) indicated strong separation of the clusters. A highly similar three cluster structure was also detected when excluding participants with a high negative impression rating (Q=0.59), and when randomly selecting half (Q=0.6) or a quarter of the sample (Q=0.61).

The cluster assignment resulted in roughly equal splits between the three clusters (Cluster 1: 150(33.93%), Cluster 2: 145(32.80%), Cluster 3: 147(33.25%). There were significant differences on all subscales of the Conners 3 questionnaire between groups (see Figure 3 and Table 3). Children in the clusters were characterised either by problems associated with cognitive control (C1: Inattention, Hyperactivity/Impulsivity/Executive Function), learning difficulties (C2: Learning Problems), or behavioural conduct problems (C3: Aggression, Peer Relations). Standardised scores indicated that the majority of children in the current sample scored in the elevated to highly elevated range across all Conners subscales relative to age-norms. The profiles based on scaled raw scores were also apparent when using the age-standardised scores. (see Figure 3b).

**Figure 3:**
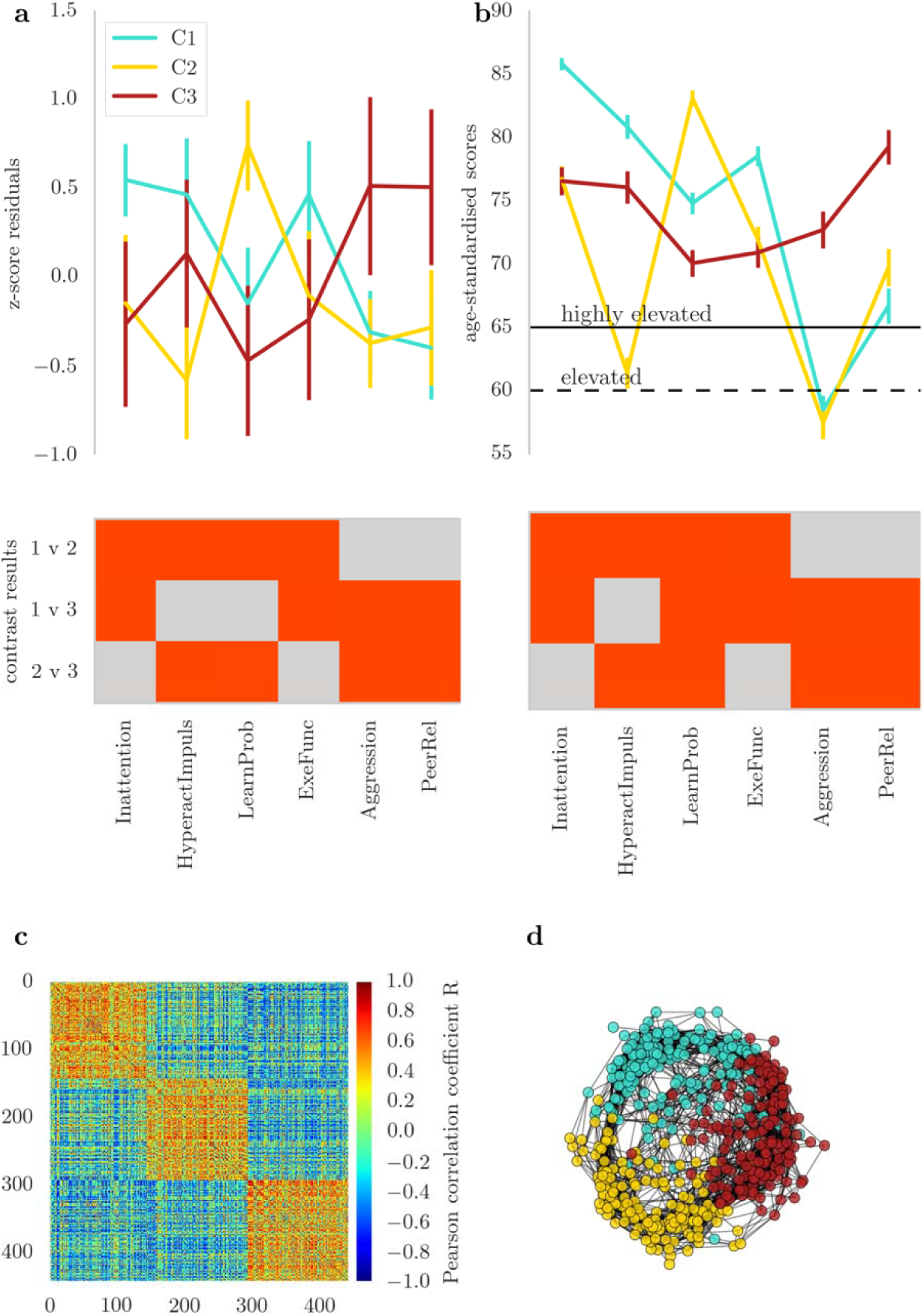
**a** Profile of ratings on the Conners 3 questionnaire in the three clusters indicated by the community detection algorithm. The top of the figure shows the mean of scores in each group with two standard errors. The scores represent residuals after regressing the effect of age. The bottom figure shows the results of group-wise contrasts on each scale.eRed indicates a significant difference between groups ( ) after Bonferroni correction. **b** Comparison of the groups on scores standardised with reference to the normative data of the Conners-3 questionnaire. **c** Child-by-child correlation matrix of Conners-3 scores after ordering the matrix according to the cluster assignment indicated by consensus clustering. The order matrix shows a clear separation between the clusters. **d** Correlation matrix in a spring layout colour-coded according to the cluster assignmen t indicated by consensus clustering. The spring layout representation shows clear spatial separation between the clusters.

**Table 3:** Scales of the Conners-3 questionnaire: Inattention, Hyperactivity/Impulsivity (HyperactImpuls.), Learning Problems (LearnProb.), Executive Function (ExeFunc), Aggression, Peer Relationship Problems (PeerRel.); mad: median absolute deviance; U: Mann Whitney U statistic; all p-values are Bonferroni corrected.

|  | Learning (1) |  | Exec. Function (2) |  | Conduct (3) |  | 1 v 2 |  | 1 v 3 |  | 2 v 3 |  |
| --- | --- | --- | --- | --- | --- | --- | --- | --- | --- | --- | --- | --- |
|  | median | mad | median | mad | median | mad | U | p | U | p | U | p |
| Inattention | 0.71 | 0.404 | 0.11 | 0.810 | 0.01 | 0.959 | 4867 | p<0.001 | 6487 | p<0.001 | 12026 | 1.000 |
| HyperactImpuls. | 0.63 | 0.636 | -0.76 | 0.682 | 0.38 | 0.886 | 3149 | p<0.001 | 9294 | 0.133 | 7321 | p<0.001 |
| LearnProb. | -0.15 | 0.642 | 0.90 | 0.524 | -0.58 | 0.876 | 3269 | p<0.001 | 9051 | 0.052 | 4137 | p<0.001 |
| ExeFunc | 0.60 | 0.599 | -0.10 | 0.760 | -0.13 | 0.953 | 5427 | p<0.001 | 7027 | p<0.001 | 11782 | 1.000 |
| Aggression | -0.53 | 0.452 | -0.59 | 0.497 | 0.26 | 1.048 | 7649 | 0.843 | 6885 | p<0.001 | 6871 | p<0.001 |
| PeerRel. | -0.56 | 0.571 | -0.45 | 0.665 | 0.67 | 0.938 | 8085 | 1.000 | 5744 | p<0.001 | 7099 | p<0.001 |

Next, the prevalence of pre-existing diagnoses in each cluster was evaluated. Children with a diagnosis of ADHD were over-represented in C1: Inattention, Hyperactivity/Impulsivity/Executive Function (see Table 4 for a breakdown of diagnoses per cluster, *X*^2^(3,354)=72.87, *p*=0.000). Other diagnoses were equally distributed between the clusters (ASD: *X*^2^(3,354)=0.06, *p*=0.971, Anxiety/Depression: *X*^2^(3,354)=0.54, *p*=0.764, Learning Deficit: *X*^2^(3,354)=3.88, *p*=0.144).

**Table 4:** Breakdown of diagnoses in each cluster identified through data-driven clustering.

|  | C1 | C2 | C3 | Total |
| --- | --- | --- | --- | --- |
| ADHD | 33 | 4 | 24 | 61 |
| ASD | 13 | 5 | 6 | 24 |
| Learning Deficit | 3 | 22 | 7 | 32 |
| Other | 7 | 8 | 8 | 23 |
| None | 94 | 106 | 102 | 302 |
| Total | 150 | 145 | 147 | 442 |

## Subgroups show differences in other questionnaire measures of executive function and everyday difficulties

Next, the groups defined through community assignment based on Conners-3 data were compared on other questionnaire measures of behavioural problems linked to executive function difficulties (BRIEF) and everyday behavioural problems (SDQ). A comparison of these measures indicated significant differences between the groups. For the BRIEF, children in Cluster 1 (Inattention/hyperactivity/Executive Function) had more problems with working memory. Children in Cluster 2 (Learning Problems) were rated as having fewer difficulties with inhibition and monitoring and Cluster 3 (Aggression/Peer Problems) were also rated as having significantly higher problems in emotional control compared to the other groups (see Figure 4a).

**Figure 4:**
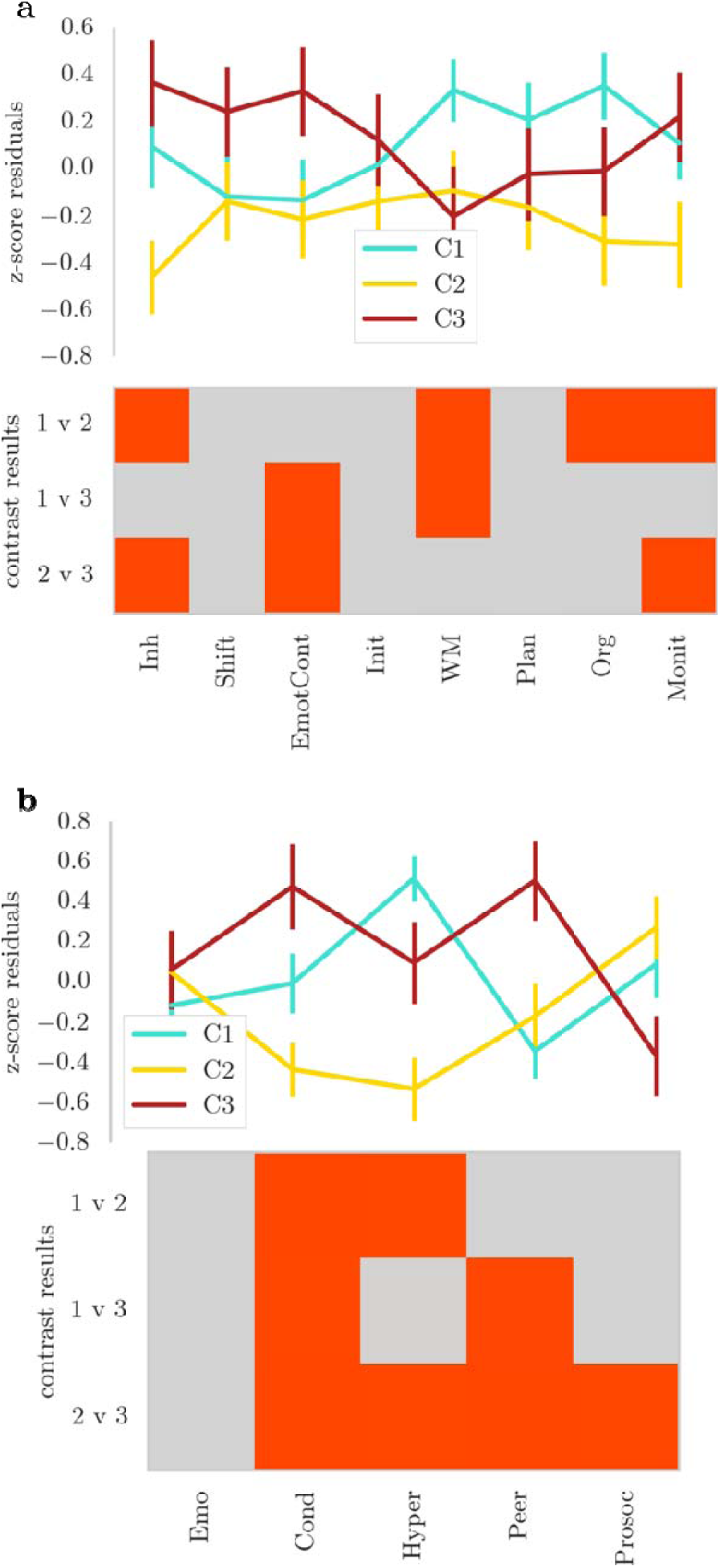
Profile of ratings for children in the clusters defined by consensus module assignment on **a**) a questionnaire on executive function difficulties (BRIEF) and **b**) a questionnaire on strengths and difficulties (SDQ). The lines indicate the mean of each group across the questionnaire scales with error bars showing two standard errors around the mean. The bottom of each figure shows the binary outcome of t-tests comparing the groups. Red indicates a significant result ( ) after Bonferroni correction. Please note that higher scores indicate a higher level of difficulties on each scale, apart from the Prosocial Behaviour (Prosoc) scale where high scores indicate more prosocial behaviour. Abbreviations: top: Inh=Inhibition, EmotCont=Emotional Control, Init=Initiate, WM=Wokring Memory, Org=Organisation of Materials, Monit=Monitoring;g; bottom: Emo=Emotional Problems, Cond=Conduct Problems, Hyper=Hyperactivity, Peer=Peer Problems, Prosoc=Prosocial Behaviour.

For the strengths and difficulties questionnaire, children in Cluster 1 (Inattention/hyperactivity/Executive Function) were characterised by high ratings for hyperactivity compared to Cluster 2 (Learning Problems), but lower conduct and peer relationship problem ratings compared to Cluster 3 (Aggression/Peer Problems). Children in Cluster 2 (learning problems) received significantly lower ratings for problems related to hyperactivity. Children in Cluster 3 (Aggression/Peer Problems) received significantly higher ratings for conduct and peer relationship problems (see Figure 4b).

## Data-driven grouping leads to more homogeneous behavioural profiles

The novel recruitment method of our sample, which includes children with specific, multiple and no diagnosis, enabled us to explore the homogeneity of the behavioural profiles within established diagnostic categories by comparison with our data-driven groupings. For the statistical comparison, a random sample of 15 (65% of the smallest sample size) was drawn from all participants within a group and correlations between their scales were calculated. This was repeated 1000 times to create a bootstrap sample of correlations. The correlations were averaged over the three data-driven groups and over the four diagnostic categories (ADHD, ASD, Learning Deficit, Other). The statistical comparison indicated that the difference between correlations in the data-driven groups and the diagnostic groups was significantly above 0 indicating higher correlations in the data-driven grouping (n=1,000, mean=0.23, SE=0.001, p=0.001). Crucially, the similarity was also significantly higher when comparing the data-driven groups to diagnostic groups on other questionnaires that were not used to inform the clustering algorithm (BRIEF: mean=0.15, SE=0.001, p=0.001; SDQ: mean=0.07, SE=0.001, p=0.026). This indicates that the data-driven grouping has identified groups of children with more common profiles of behavioural symptomatology than we would expect to find in children grouped on the basis of more traditional diagnostic criteria.

## Subgroups show differences in the structural connectome

Next, we investigated the relationship between white matter connectivity and the groups defined through consensus clustering using partial least squares (PLS) regression. The first three PLS components explained 48% of variance in group membership (Component 1: 21.23% (SD: 4.302); Component 2: 16.28% (SD: 5.944); Component 3: 10.57% (SD: 4.277), bootstrapped mean and standard deviation (SD) over 1000 permutations). Further components explained less than 5% of variance and were therefore dropped from the analysis. Comparison of component loadings per group indicated significant lower loading of C1 (Inattention/Hyperactivity/Executive Function) compared to the other groups for PLS component 1, significantly higher loading in C1 (Inattention/Hyperactivity/Executive Function) compared to C3 (Aggression/Peer Problems) for PLS component 2, and significantly lower loading in C1 (Inattention/ Hyperactivity/Executive Function) compared to C2 (Learning Problems) for PLS component 3.

There were differences in the brain areas that distinguished the groups. PLS 1 that distinguished between C1 (Inattention/Hyperactivity/Executive Function) and the other groups loaded most heavily on the rostral middle frontal, superior frontal, lateral orbitofrontal, anterior cingulate, lateral occipital and fusiform cortex (see Figure 5). The second PLS component, which distinguished between C2 (Learning Problems) and C1 (Inattention/Hyperactivity/Executive Function), loaded the most on the rostral middle frontal, lateral orbitofrontal, anterior and posterior cingulate, and lateral occipital cortex. The third PLS component, which distinguished C3 (Aggression/Peer Problems) from the other groups, loaded on the lateral orbitofrontal, anterior cingulate, and entorhinal cortex, and also on connections of the right pallidum and putamen (see Table 5).

**Table 5:** PLS scores for subcortical areas

|  |  | PLS 1 | PLS 2 | PLS 3 |
| --- | --- | --- | --- | --- |
| left | accumbens | 0 | 0 | 0 |
|  | amygdala | 0 | 0 | 0 |
|  | caudate | 0 | 0 | 0 |
|  | hippocampus | 0 | 0 | 0 |
|  | pallidum | 0 | 0 | 0 |
|  | putamen | 0 | 0 | 0 |
|  | thalamus | 0 | 0 | 0 |
| right | accumbens | 0 | 0 | 0 |
|  | amygdala | 0 | 0 | 0 |
|  | caudate | 0 | 0 | 0 |
|  | hippocampus | 0 | 0 | 0 |
|  | pallidum | 0 | 0 | 35 |
|  | putamen | 0 | 0 | 37 |
|  | thalamus | 0 | 0 | 0 |

**Figure 5:**
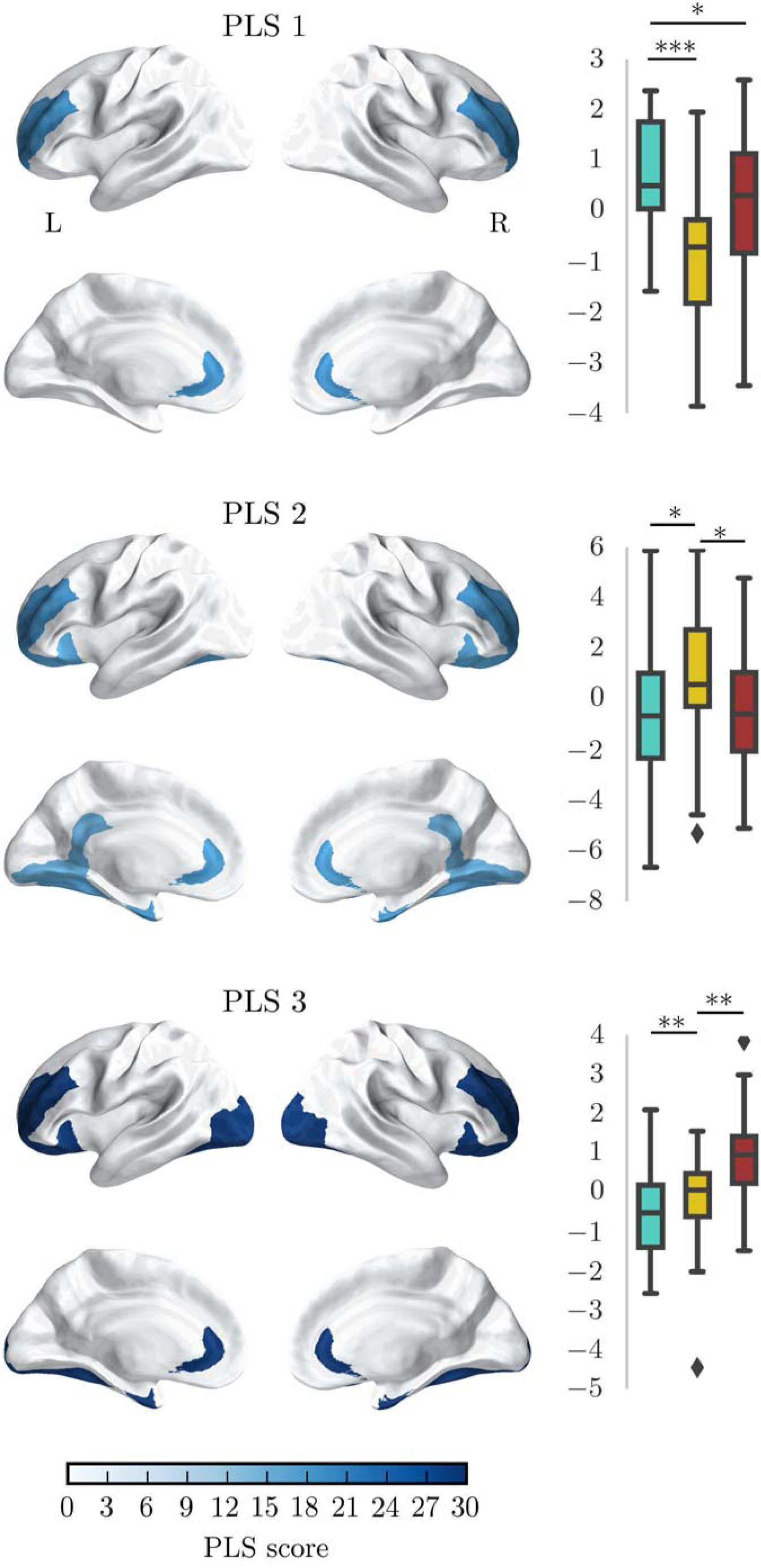
Relationship between the node degree of brain regions in the structural connectome and clusters based on Conners-3 responses. The brain maps show the score of PLS components for brain regions that most strongly distinguished the group (top 20%). PLS scores above 2 are considered to be significantly predictive. The graphs show the statistical comparison of groups on loadings for each component.

## Discussion

In the present study, we use a data-driven clustering algorithm to group children according to their similarity on ratings of behaviours associated with attention problems. The findings demonstrated that among a large sample of children referred by clinical and educational specialists for cognitive learning problems there exist distinct behavioural profiles that transcend traditional diagnostic categories. Three groups were identified: one with problems related to executive function, inattention and hyperactivity, a second with severe learning difficulties, and a third with behavioural conduct problems. These groups were consistent in randomly-selected subsets of the sample, and reliably reproduced in simulated data with a known structure, even when adding considerable noise. The three behavioural profiles identified were evident in two additional parent rating scales that were not used to inform the original algorithm. Further, comparison of white matter connectivity indicated that the data-driven groups were distinguished by connectivity of the lateral prefrontal and cingulate cortex

One of the subgroups was characterised by elevated symptoms of inattention, hyperactivity/impulsivity, and executive function. This group was also rated as having increased difficulties with behaviours relating to working memory, organisation, planning, and hyperactivity on two other rating scales that were not used as part of the clustering algorithm. This behavioural profile captures core problems associated with the traditional ADHD diagnostic label, which too is marked by high levels of inattention and hyperactivity and executive function problems {Barkley:1997vr, Castellanos:2002hq, Pennington:1996fq, ^36^. A disproportionate number of children with an ADHD diagnosis were assigned to this cluster. However, this subtype was not synonymous with ADHD as half of the children with an ADHD diagnoses were split across the other two clusters that were defined by markedly different behavioural profiles.

A second subgroup had severe learning deficits relative to the other two groups. On other questionnaires, they were rated as having fewer problems with inhibition, attention and other aspects of executive function compared to children in the other clusters. However, their scores on these scales were in the elevated and clinical range when compared to age-norms, indicating that they fall below age expected levels for attention and executive function, but have less pronounced difficulties in these areas than children in the other clusters. Overall, this group displayed elevated symptoms of inattention and executive function difficulties combined with fewer problems with hyperactivity/impulsivity. This profile resembles that described for the inattentive subtype of ADHD^37^, but it should be noted that the most distinguishing feature of this group was pronounced learning difficulties.

A third subgroup was characterised by difficulties with aggression and peer relationships. Children in this group were also rated as having increased problems with behaviours related to emotional control and conduct on the two rating scales not used as part of the clustering algorithm. The distinction between groups with problems relating to either executive function or behavioural conduct is reminiscent of the debate surrounding the overlap between ADHD and oppositional defiant disorder (ODD)/conduct disorder (CD). Some authors have argued for a high degree of overlap between these diagnostic groups ^4^, but evidence from genetic and imaging studies had suggested distinct pathophysiological mechanisms ^38-40^. Consistent with these results, the current study shows that behavioural ratings of inattention/hyperactivity and aggression/peer relationship problems form distinct clusters.

These results demonstrate that data-driven clustering using a community detection algorithm can be used to characterise common and complex behavioural problems in children. The clustering algorithm identified groups that mirror some of the distinction of traditional diagnostic groups. One of the major advantages of this approach is that more homogeneous groupings were identified, and these are better suited for investigations into underlying biological mechanisms. While not the main aim of the current analysis, exploratory analysis showed that our data-driven sub-grouping was associated with underlying differences in structural connectivity between groups. The areas distinguishing the groups have been suggested to play a role in relevant behaviours, making it possible to formulate hypotheses about neurobiological mechanisms associated with the different behavioural profiles. For instance, the group characterised by problems relating to attention and executive function showed differences in connectivity of the prefrontal, anterior cingulate cortex, and lateral occipital cortex. These differences in white matter connections of circuits related to inhibitory control^41^, goal-directed behaviour and visual attention ^43^ may play a role in the aetiology of these behavioural problems. In contrast, children with a profile of problems relating to emotional regulation and peer relationships were distinguished from the other groups by differences in white matter connectivity of the rostrolateral prefrontal cortex, anterior cingulate cortex, pallidum, and putamen. These findings may imply a difference in integration between the prefrontal cortex and the basal ganglia system {Finger:2011kd, ^44^.

In summary, clustering of similarities across behavioural problem identified three groups with distinct profiles of difficulties that related to inattention, learning, and peer relationships respectively. These groups were also distinguished by the connectivity of circuits previously implicated in executive function and behavioural regulation, including the prefrontal cortex, cingulate cortex, and their subcortical connections. These findings act as an important proof of principle: data-driven profiling provides an alternative means of distinguishing common and complex behavioural problems in children. This data-driven classification of behavioural problems may provide a better account of behavioural differences and relate more closely to neurobiological mechanisms than traditional diagnostic approaches.

## Acknowledgements

The Centre for Attention Learning and Memory (CALM) research clinic is based at and supported by funding from the MRC Cognition and Brain Sciences Unit, University of Cambridge. The Principal Investigators are Joni Holmes (Head of CALM), Susan Gathercole (Chair of CALM Management Committee), Duncan Astle, Tom Manly and Rogier Kievit. Data collection is assisted by a team of researchers and PhD students at the CBSU that includes Sarah Bishop, Annie Bryant, Sally Butterfield, Fanchea Daily, Laura Forde, Erin Hawkins, Sinead O’Brien, Cliodhna O’Leary, Joseph Rennie, and Mengya Zhang. The authors wish to thank the many professionals working in children’s services in the South-East and East of England for their support, and to the children and their families for giving up their time to visit the clinic.

## Supplementary Analyses

### Robustness of the consensus clustering algorithm

In order to test the reliability of the community detection algorithm under varying conditions, random networks with known community structure were created. The networks consisted of 100 nodes with 4 modules. The connection likelihood within and between clusters was systematically varied between 0.1 and 0.9. The quality index of the community structure was calculated at each combination of between-and within-cluster connection likelihood. The results indicated a high-quality index for network with higher within-cluster than outside-cluster connection likelihood (see Figure S1a). High connection density outside of clusters had a large influence, even when the connection likelihood within modules was very high.

For comparison with the empirical network of Conners-3 score correlations, the connection density within and between networks was calculated. To this end, all connections were binarized so that any connection with a Pearson correlation coefficient above 0 was set to 1. The connection density was estimated as the ratio between existing connections in the binarized empirical network and a fully connected network of the same size. Connection density within modules based on consensus clustering was 0.79 and connection density between modules was 0.05. Together with the results of the simulated networks, these connection densities indicate very high separation of the network clusters.

We further tested the robustness of the community assignment by adding increasing percentages of random Gaussian noise (*μ*=0, *σ*=1) to the network matrix and repeated the consensus clustering procedure (see Figure S1b). The quality index indicated good separation of the clusters between 5 and 30% noise (Q between 0.62 and 0.65). No stable assignment could be reached at 35% of noise and above. These results indicate that the community assignment is robust to a considerable amount of noise.

**Figure S1:**
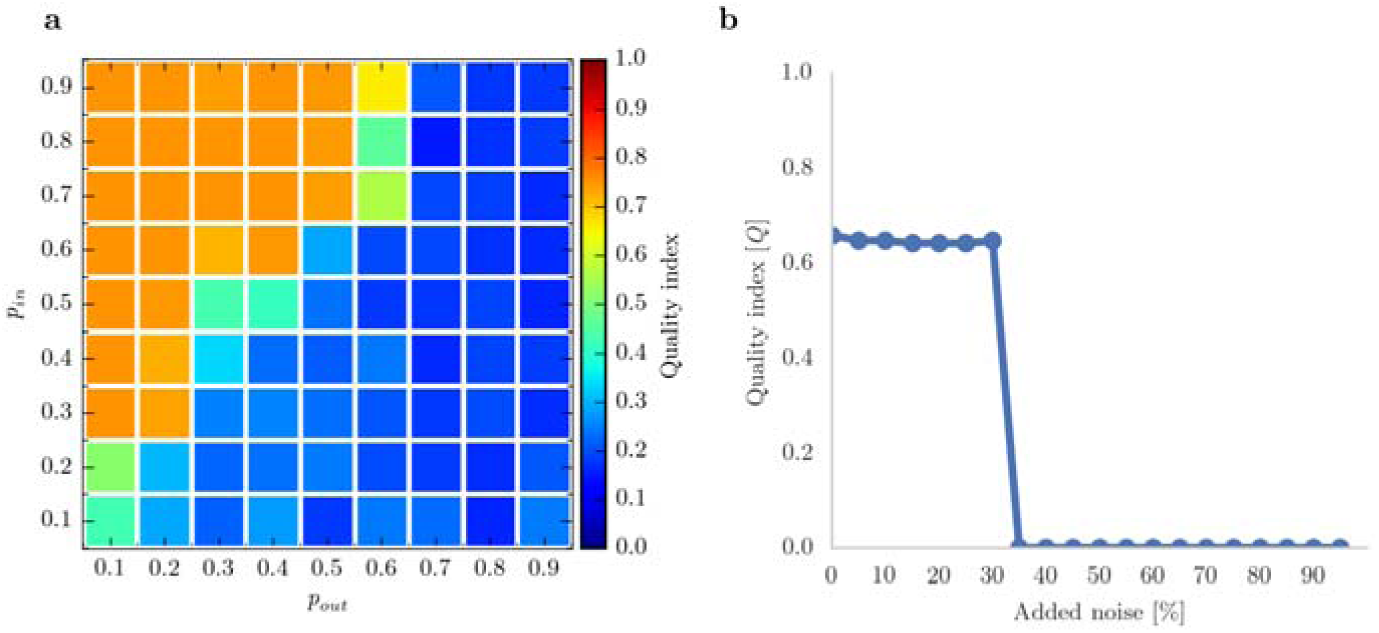
Results of robustness testing **a**: Quality indices of consensus clustering using simulated networks with varying levels of within ( ) and between ( ) connections probabilities. High within-cluster and low between-cluster connectivity lead to high separation of clusters with consensus clustering, i.e. high quality-indices. **b**: Consensus clustering using the empirical child-by-child network of Conners-3 correlations with varying levels of added noise. The three-cluster solution could be reconstructed up to 30% of added Gaussian noise. At higher-level of noise, no clustering solution could be obtained.

### Influence of connection weight and thresholding on structural connectome results

Different methods exist for the construction of structural networks from diffusion-weighted data and there is currently no scientific consensus on the best approach (Qi, Meesters, Nicolay, ter Haar Romeny, & Ossenblok, 2015). Networks in the current analysiss were waited by fractional anisotropy (FA), a commonly used measure of white matter organisation based on the diffusion tensor model. FA characterises the directedness of diffusion within a voxel, but may lead to misinterpretation in regions of crossing fibres (Douaud et al., 2011). Therefore, the main analysis was repeated with networks weighted by Generalized FA (GFA) based on a constant solid angle (CSA) model, which is better able to take crossing fibres into account (Tuch, 2004). The node degree for each brain region was identical for the GFA and FA model for density thresholds between 5% and 15% (Kolmogorov-Smirnov test for two samples: p=1.0 uncorrected for all regions). It follows that the PLS analysis provides the same results for networks weighted by FA and GFA as this analysis was based on node degrees and node degrees were identical for both models within the relevant density range.

Another potential source of variation in the analysis is the density threshold. Network analyses are sensitive to the number of connections. Therefore, density thresholding is often applied, but the chosen threshold may influence the results of the analysis. For the current investigation, the influence of different density thresholds was systematically investigated by repeating the analysis over a range of densities and comparing the factor scores in a repeated-measures analysis of variance model with factors for density and the interaction between density and component (components loading density + component + density*component). The results indicated no significant effect of density or the interaction between density and any component (model fit: F(9, 14790)0.001, *p*=1, Adjusted-*R^2^*=-0.001; Density: *t* <0.001, *p*=1, Interactions: *t* <0.001, *p*=1).

